# Time dependent effects of cerebellar tDCS on cerebello-cortical connectivity networks in young adults

**DOI:** 10.1101/2023.06.26.546626

**Authors:** Ted Maldonado, T. Bryan Jackson, Jessica A. Bernard

## Abstract

The cerebellum is involved in non-motor processing, supported by topographically distinct cerebellar activations and closed loop circuits between the cerebellum and the cortex. Disruptions to cerebellar function and network connectivity in aging or disease may negatively impact prefrontal function and processing. Cerebellar resources may be important for offloading cortical processing, providing crucial scaffolding for normative performance and function. Here, we used transcranial direct current stimulation (tDCS) to temporarily alter cerebellar function and subsequently investigated resting state network connectivity. This allows us to investigate network changes that may parallel what is seen in aging and clinical populations, providing additional insights into these key circuits. Critically, what happens to these circuits if the cerebellum is not functioning optimally remains relatively unknown. We employed a between-subjects design applying anodal (n=25), cathodal (n=25), or sham (n=24) stimulation to the cerebellum to examine the effect of stimulation on cerebello-cortical resting state connectivity in young adults. We predicted increased functional connectivity following cathodal stimulation and decreased functional connectivity following anodal stimulation. We found, anodal stimulation resulted in increased connectivity in both ipsilateral and contralateral regions of the cortex, perhaps indicative of a compensatory response to degraded cerebellar output. Additionally, a sliding window analysis also demonstrated a time dependent nature to the impacts of cerebellar tDCS on connectivity, particularly in cognitive region in the cortex. Assuming the difference in connectivity and network-behavior relationships here parallels what occurs in aging or disease, this may provide a mechanism whereby offloading of function to the cerebellum is negatively impacted, resulting in subsequent differences in prefrontal cortical activation patterns and performance deficits. These results might inform and update existing compensatory models of function to include the cerebellum as a vital structure needed for scaffolding.

## Introduction

Viral tract tracing in non-human primates has shown topographically distinct loops connecting the cerebellum and cortex. Anterior regions such as lobules I-IV and V have projections to the primary motor cortex, whereas posterior lobules such as Crus II project to the prefrontal cortex (Kelly & Strick, 2003). Converging evidence in human neuroimaging parallels the viral tract tracing work in primates (Clower et al., 2005; K. A. Coffman et al., 2011; Hoshi et al., 2005; Kelly & Strick, 2003). Both human functional magnetic resonance imaging (fcMRI; Krienen & Buckner, 2009) and diffusion-weighted imaging (Salmi et al., 2010) found segregated fronto-cerebellar networks, linking the cerebellum to the motor cortex, the dorsal lateral prefrontal cortex, the medial prefrontal cortex, and the anterior prefrontal cortex in humans. O’Reilly and colleagues (2010) have furthered this understanding by demonstrating functional zones in the cerebellum such that anterior regions of the cerebellum have specific projections to motor, somatosensory, visual, and auditory cortices; whereas, posterior lobes have projections to, and functional networks with, the prefrontal and posterior parietal cortices (Bernard et al., 2012; Diedrichsen et al., 2019; Guell et al., 2018; King et al., 2019; O’Reilly et al., 2010). Behaviorally, work has suggested connectivity between the PFC and cerebellum might predict performance on learning executive function tasks (Reineberg et al., 2015), highlighting the non-motor contributions of the structure and its networks.

Critically, little work has examined how differences and changes in these functional networks might affect cognitive performance and communication between brain regions, though work in aging (Bernard et al., 2013) and disease (Allen et al., 2007; Bai et al., 2009; Bernard & Mittal, 2014; Mittal et al., 2014) might begin to provide clues. Degraded gray and white matter integrity in aging and disease have behavioral implications as the internal models used by the cerebellum for automaticity may not be utilized as effectively due to disruptions in cerebellar function and network connectivity (Bernard et al., 2013; Bernard & Seidler, 2014; Maldonado, et al., 2022; Miller et al., 2013). Additionally, work suggests the cerebellum might support cortical processing, such that degraded cerebellar output results in poor task performance (Bernard, 2022; Bernard et al., 2020; Bernard & Seidler, 2014; Schmahmann et al., 2019). Put simply, automatic processing relies on internal models and cerebellar processing, freeing up cortical resources, particularly if tasks become increasingly complicated. Thus, understanding these networks and alterations therein could be the basis for an improved understanding of cerebellar contributions to behavior.

One commonly used method of neuromodulation is transcranial direct current stimulation (tDCS). tDCS typically increases (anodal) or decreases (cathodal) neural activity in the cerebral cortex using a small amount of electrical current, impacting cortical excitability (Brunoni et al., 2012; Nitsche & Paulus, 2000; Priori, Berardelli, Rona, Accornero, & Manfredi, 1998), and in turn behavior (Coffman et al., 2014). However, anodal stimulation to the cerebellum excites inhibitory Purkinje cells, which ultimately decreases signal to the cortex (Ghez, 1991; Grimaldi et al., 2016). Cathodal stimulation to the cerebellum inhibits the same cells, resulting in increased signal to the cortex. Cerebellar tDCS has the potential to be particularly informative, as one may be able to increase or decrease the connectivity between brain regions, providing insight into how the cerebellum communicates with the cortex across modulation conditions. The impact of anodal cerebellar tDCS is of particular interest, as it may provide insights into cerebello-cortical interactions that mimic those seen in a variety of neurological and psychiatric illnesses where the cerebellum is impacted (Ferrucci et al., 2016).

Limited work has examined the effects of cerebellar tDCS on cerebello-cortical connectivity, and in this limited literature, the primary focus has been on language processing networks. Specifically, anodal stimulation increases functional connectivity from the cerebellum to cortical areas involved in the motor control of speech (Turkeltaub et al., 2016), language and speech motor regions in the left hemisphere (D’Mello et al., 2017; Turkeltaub et al., 2016), and spelling (Sebastian et al., 2017). Additionally, anodal stimulation reduced cerebellar ataxia symptoms and improved cerebellar output in individuals with cerebellar ataxia (Benussi et al., 2017). Further, anodal stimulation to the cortex also increases cortico-cerebellar connectivity when applied to the dorsal lateral prefrontal cortex (Abellaneda-Pérez et al., 2020), motor cortex (Cummiford et al., 2016) and the right posterior parietal cortex (Callan et al., 2016).

Thus, cerebello-cortical connectivity has been well-documented, and work focused on language networks has indicated that these networks are subject to modulation via cerebellar tDCS (D’Mello et al., 2017; Turkeltaub et al., 2016). Critically, recent functional imaging work, coupled with tDCS, demonstrated that disrupting the cerebellum resulted in greater cortical processing, suggesting a cerebellar scaffolding mechanism (Maldonado, et al., 2022). Here, we were interested in understanding how connectivity between the cerebellum and the cortex might support this mechanism. That is, if the cerebellum is disrupted, will connectivity increase to help maintain task performance (Bernard, 2022)? To this end, we used tDCS and fcMRI to better understand cerebello-cortical functional connectivity. This work stands to provide important new insights into the impact of cerebellar tDCS on cerebellar connectivity, and in turn potential impacts on behavior, as well as critical insights into cerebellar networks when function is altered. This latter point may in turn influence our understanding of cerebello-cortical connectivity in aging, as well as neurological or psychiatric illness. To this end, participants were randomly placed in one of three stimulation conditions (anodal, cathodal, or sham) and stimulation was applied to the right cerebellum. We used a lobular approach to our analyses (Bernard et al., 2012; Grami et al., 2021). In an effort to limit the number of comparisons and in turn false positives, we focused on hemispheric Lobules I-X as these represent areas associated with both motor and prefrontal cortical regions (Bernard et al., 2012; Buckner et al., 2011; King et al., 2019). Though past work has suggested anodal stimulation increases connectivity between the cerebellum and language networks in the cortex specifically (D’Mello et al., 2017; Turkeltaub et al., 2016), we predict that cathodal stimulation will increase connectivity whereas anodal stimulation will decrease connectivity, consistent with a compensatory response. This would be consistent with past work suggesting cerebellar tDCS manipulates the inhibitory nature of Purkinjie cells (Galea et al., 2009; Ghez, 1991; Grimaldi et al., 2016), which might affect patterns in cortical activation (Maldonado et al., 2022).

## Methods

### Participants

Seventy-five healthy, young adults participated in this study and were provided monetary compensation for their time. Exclusion criteria included left handedness, history of neurological or mood disorders, skin conditions, pregnancy, and history of concussion. Data from one participant was not used because the participant did not wish to finish the experiment after providing consent. Thus, seventy-four right-handed participants (38 female) ages 18 to 30 (*M*= 22.0 years, *SD*= 3.45) were included in the analyses, a sample in line with typical imaging studies (Szucs & Ioannidis, 2020). Participants were randomly assigned to either the anodal (n=25), cathodal (n=25), or sham (n=24) stimulation condition. All procedures completed by participants were approved by the Texas A&M University Institutional Review Board and conducted according to the principles expressed in the Declaration of Helsinki.

### Procedure

Resting state data was collected as part of a larger imaging study which took approximately two hours (Maldonado, Bryan Jackson, et al., 2022). In the context of this larger study, the resting state data was collected first, immediately after entering the scanner and all data were collected within 45 minutes of the stimulation. Following the completion of the consent form, participants completed a basic demographic survey, followed by tDCS (see below for details). Participants were blind to the stimulation condition and were tDCS naïve. Following stimulation, participants were escorted to the scanner for the brain imaging protocol. Once the scanning procedures were complete, participants completed a tDCS sensation survey and were debriefed. No adverse events were reported in the tDCS sensation survey.

### tDCS Stimulation Parameters

Participants were fitted with a classic two electrode montage to administer either cathodal, anodal, or sham stimulation using a Soterix 1x1 tES system. One 5x5cm electrode was placed two cm below and four cm lateral of the inion over the right cerebellum (Ferrucci et al., 2015). The second 5x5cm electrode was placed on the right deltoid. Both electrodes were affixed using elastic bands. Each electrode was placed in a pre-soaked sponge, which had 6 mL of saline solution added on each side.

Once electrodes were placed, stimulation was set to 1.0 mA for thirty seconds to ensure the electrodes made a good connection with the scalp. If contact quality was below 50%, adjustments, such as moving hair to increase the electrode’s contact with the scalp, were made and contact quality was rechecked. Following a successful re-check, participants completed a 20-minute stimulation session at 2 mA (Ferrucci et al., 2015; Grimaldi et al., 2014, 2016). During the stimulation conditions, maximum stimulation intensity was reached in 30 seconds and maintained for 20 minutes, and then would return to 0 mA. During sham conditions, maximum stimulation intensity would be reached, but immediately return to 0 mA. There was no additional stimulation during the 20-minute sham session.

### fMRI data acquisition

Resting state fMRI data was collected at the Texas A&M Translational Imaging Center with a 3-T Siemens Magnetom Verio scanner using a 32-channel head coil. Two blood oxygen level dependent (BOLD) whole brain scans with a multiband factor of 4 were collected in the absence of any task (number of volumes = 114, repetition time [TR] = 2000 ms, echo time [TE] = 27 ms; flip angle [FA] = 52°, 3.0 × 3.0 × 3.0 mm3 voxels; 56 slices, interleaved, slice thickness=3.00mm, field of view (FOV) = 300 × 300 mm; time = 4:00 min). The scans were collected in opposite encoding directions (anterior → posterior and posterior → anterior). During the scan participants viewed a centrally located fixation cross and were asked to stay awake while “thinking about nothing in particular”. An additional high resolution T1 weighted whole brain anatomical scan was taken (sagittal; GRAPPA with acceleration factor of 2; TR = 2400 ms; TE = 2.07 ms; 0.8 x 0.8 x 0.8 mm3 voxels; 56 slices, interleaved, slice thickness= 0.8mm; FOV = 256 × 256 mm; FA = 8°; time = 7:02 min) for data normalization.

### Rs-fMRI data pre-processing

Both anatomical and functional images were collected using DICOM format but were converted to NIFTI files and organized into a Brain Imaging Data Structure (BIDS) using bidskit (v 2019.8.16; Mike Tyszak, 2016).

Rs-fMRI data were pre-processed using CONN (Whitfield-Gabrieli & Nieto-Castanon, 2012) standalone toolbox (v20b). Functional and anatomical data were preprocessed using a flexible preprocessing pipeline (Nieto-Castanon, 2020b) including realignment with correction of susceptibility distortion interactions, slice timing correction, outlier detection, direct segmentation and MNI-space normalization, and smoothing. Functional data were realigned using SPM realign & unwarp procedure (Andersson et al., 2001), where all scans were coregistered to a reference image (first scan of the first session) using a least squares approach and a 6 parameter (rigid body) transformation (Friston et al., 1995), and resampled using b-spline interpolation. Temporal misalignment between different slices of the functional data (acquired in interleaved Siemens order) was corrected following SPM slice-timing correction (STC) procedure (Henson et al., 1999; Sladky et al., 2011), using sinc temporal interpolation to resample each slice BOLD timeseries to a common mid-acquisition time. Potential outlier scans were identified using ART (Whitfield-Gabrieli et al., 2011) as acquisitions with framewise displacement above 0.9 mm or global BOLD signal changes above 5 standard deviations (Nieto-Castanon, 2022; Power et al., 2014), and a reference BOLD image was computed for each subject by averaging all scans excluding outliers. Functional and anatomical data were normalized into standard MNI space, segmented into grey matter, white matter, and CSF tissue classes, and resampled to 2 mm isotropic voxels following a direct normalization procedure (Calhoun et al., 2017; Nieto-Castanon, 2022) using SPM unified segmentation and normalization algorithm (Ashburner, 2007; Ashburner & Friston, 2005) with the default IXI-549 tissue probability map template. Last, functional data were smoothed using spatial convolution with a Gaussian kernel of 5 mm full width half maximum (FWHM).

### Functional Connectivity Analysis

Functional data were denoised using a standard denoising pipeline (Nieto-Castanon, 2020a) including the regression of potential confounding effects characterized by white matter timeseries (5 CompCor noise components), CSF timeseries (5 CompCor noise components), motion parameters and their first order derivatives (12 factors) (Friston et al., 1996), outlier scans (12 factors) (Power et al., 2014), and linear trends (2 factors) within each functional run, followed by bandpass frequency filtering of the BOLD timeseries (Hallquist et al., 2013) between 0.008 Hz and 0.09 Hz. This was applied to the resting state data in order to focus on slow-frequency fluctuations while minimizing the influence of physiological, head-motion and other noise sources (Hallquist et al., 2013). Potential confounding effects to the estimated BOLD signal were determined and removed separately for each voxel and for each subject and functional run/session, using Ordinary Least Squares (OLS) regression to project each BOLD signal timeseries to the sub-space orthogonal to all potential confounding effects using an anatomical component-based noise correction procedure (aCompCor) (Behzadi et al., 2007).

*Seed to voxel analysis*. Seed to voxel analyses, were conducted using *a priori* whole lobular (Bernard et al., 2012; Grami et al., 2021) ROIs in the right cerebellar hemisphere (Lobule I-IV, V, VI, VIIb, Crus I, Crus II, VIIIA, VIIIB, XI, and X). Fisher-transformed bivariate correlation coefficients between each lobular ROI BOLD timeseries and each individual voxel BOLD timeseries were assessed for the anodal, cathodal, and sham stimulation groups individually. Individual scans were weighted by a boxcar signal characterizing each individual task or experimental condition convolved with an SPM canonical hemodynamic response function and rectified. Group level comparisons were as follows: anodal>sham, cathodal>sham, and anodal>cathodal. Results were thresholded using a combination of a cluster-forming p < 0.001 voxel-level threshold, and a familywise corrected p-FDR < 0.05 cluster-size threshold (Chumbley et al., 2010).

*Sliding Window*. A sliding window analysis was used to evaluate sources of temporal variability in functional connectivity patterns. We created five-time windows that were 120 seconds long, with onsets at 0, 30, 60, 90, and 120 seconds post scan onset. Each individual window is treated as a separate condition, and weighted GLM is used to compute the corresponding condition-/time-specific measures. Variability of these measures across time is then computed. We assessed anodal, cathodal, and sham stimulation individually and the contrasts between them for each cerebellar seed region within each time window and compared changes in connectivity. Results were thresholded using a combination of a cluster-forming p < 0.001 voxel-level threshold, and a familywise corrected p-FDR < 0.05 cluster-size threshold (Chumbley et al., 2010).

## Results

Connectivity patterns following each stimulation condition are presented in the Supplementary material. Connectivity patterns were broadly consistent with canonical cerebello-cortical networks (Bernard et al., 2012; Diedrichsen et al., 2019; King et al., 2019; O’Reilly et al., 2010) for the sham (Supplemental Table 1) and active (Supplemental Table 1) stimulation groups. In brief, for lobules I-V we primarily see positive connectivity to motor regions, such as the precentral gyrus, in addition to other memory, language and visual processing regions of the cortex. Posterior regions of the cerebellum correlated with regions within the frontal, parietal, and temporal lobes of the cortex, as expected. Here, we have focused on significant group contrasts (i.e. Anodal > Cathodal) that emerge within each lobule.

*Rest*. When examining motor-oriented lobules (Lobules I-IV and V), we found lower connectivity between Lobules I-V and regions in the right pre and post central gyrus and other motor regions in the cathodal stimulation group compared to the sham group (Figure 1A; Table 1). For the cognitively oriented lobules (Lobules VI-Crus II), we found higher connectivity between Crus II and visual regions such as the intra and supracalcarine cortex, and the cuneus in the anodal stimulation group compared to cathodal (Figure 1B; Table 1). We also found lower connectivity between Lobule X and visual regions such as the Lingual Gyrus and the lateral occipital lobe, as well as cognitive regions in the superior parietal lobe in the anodal stimulation group compared to the cathodal group (Table 1). Lastly, connectivity was lower between Lobule X and visual regions such as the right intra and supracalcarine cortex, and the cuneus in the cathodal group compared to the sham group (Table 1). Together, cathodal stimulation might result in lower connectivity between motor lobules in the right cerebellum and ipsilateral motor regions of the cortex. Anodal stimulation over cognitively oriented lobules in the right cerebellum result in higher contralateral connectivity in areas related to vision processing.

**Figure 1.**
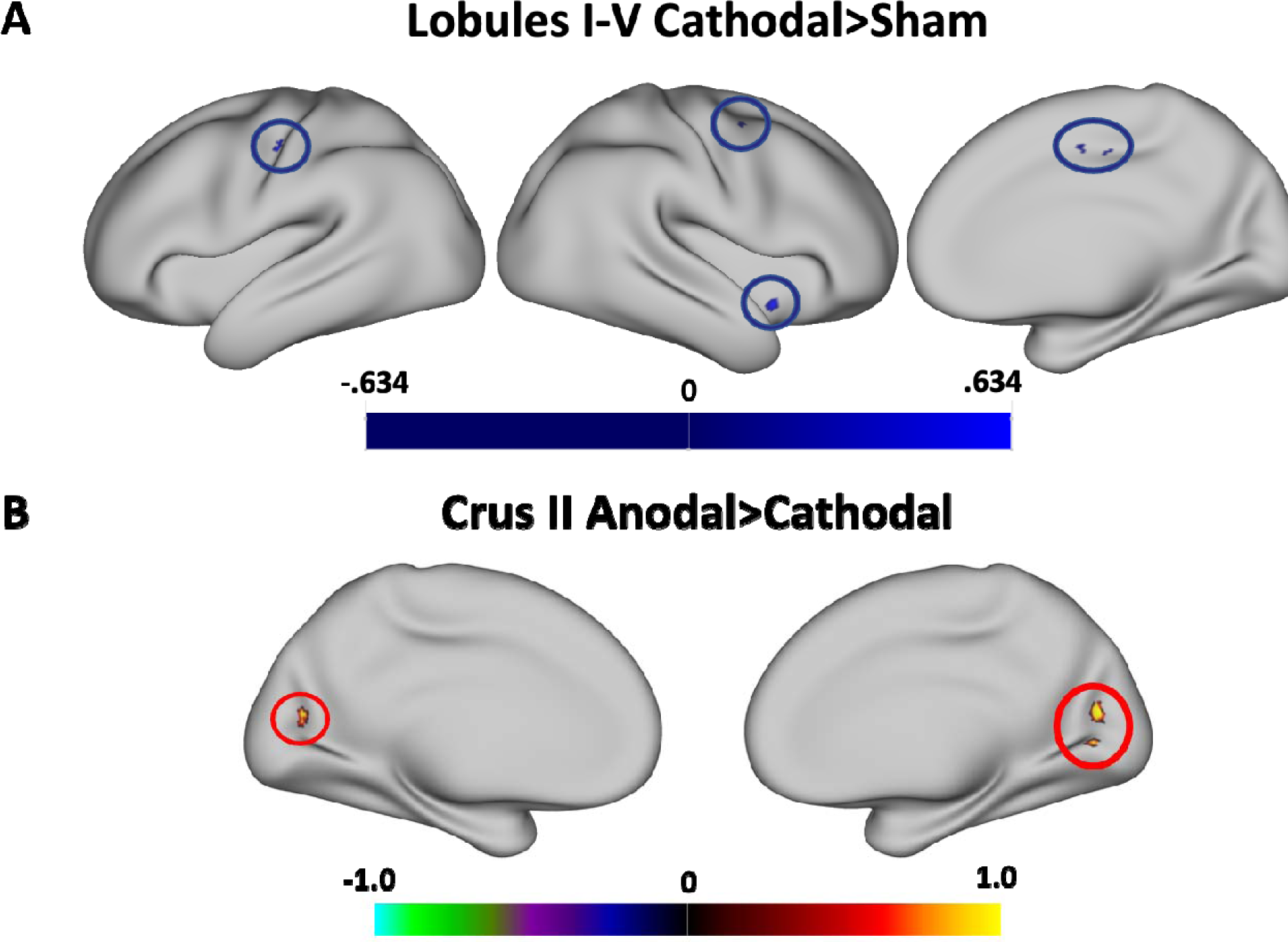
**A.** The cathodal stimulation group had lower connectivity (relative to sham) to motor regions, with some modulation in cognitively oriented regions in the frontal lobe. **B.** The anodal stimulation group showed higher connectivity (relative to cathodal) to regions in the occipital lobe. The color bars denote the percentile range of z-scores. The maps are thresholded such that only significant (p-FDR_J<_J0.05) results are presented.

**Table 1.**
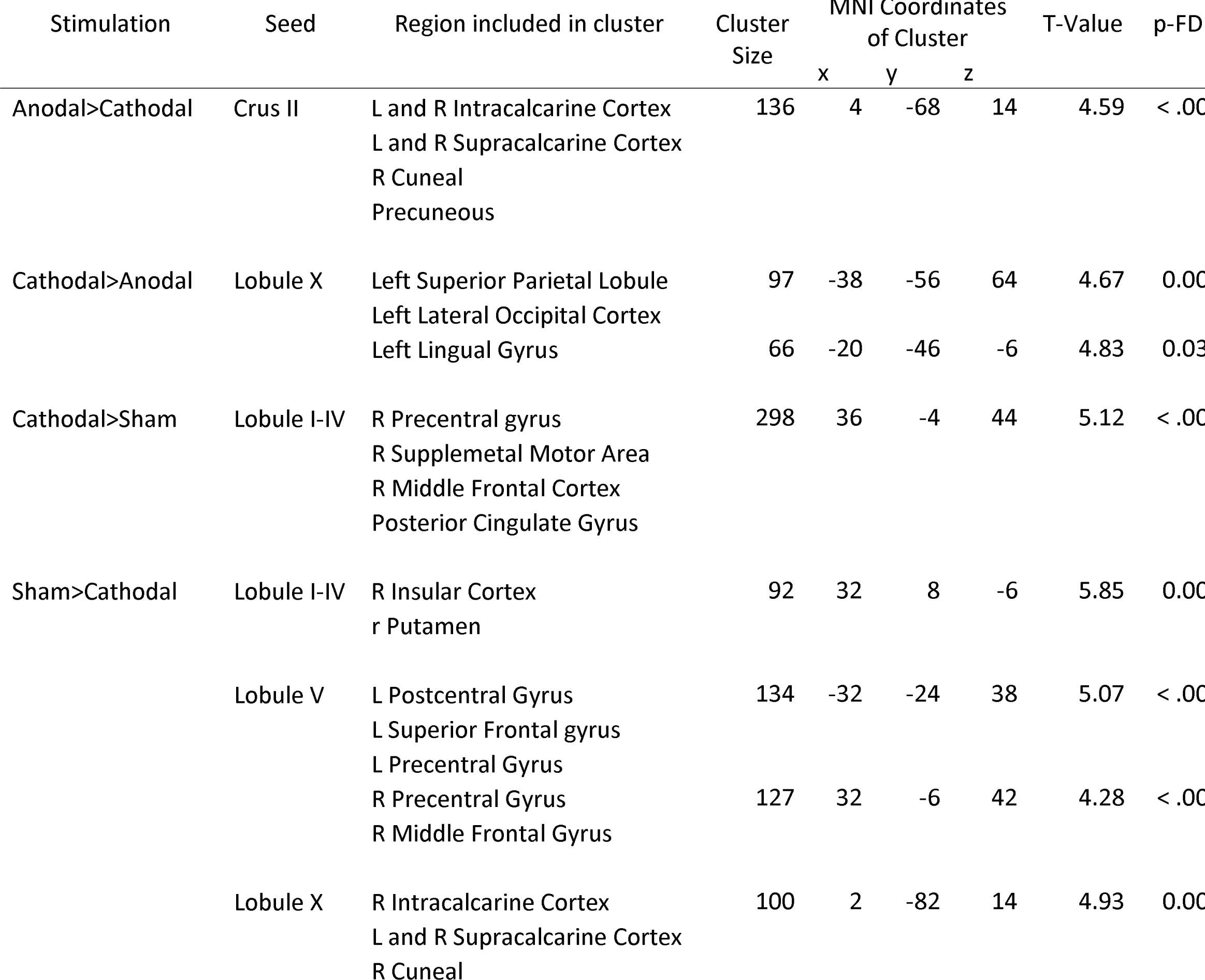
MNI coordinates of the local maxima of brain regions showing functional connectivity with the lobules of the right cerebellar hemisphere by stimulation contrast.

*Sliding window*. A sliding window analysis was used to investigate how connectivity patterns vary across the duration of the scan. Below we describe general patterns within early and late windows. Early windows were 120 seconds long, with onsets at 0 and 30 second post scan onsets, whereas late windows were also 120 seconds long, but had onsets at 60, 90, and 120 seconds.

When examining connectivity of motor lobules in the cerebellum, there was a general trend in which stimulation in the cathodal group lowered variability in connectivity from Lobule I-V to ipsilateral motor areas, such as the precentral gyrus, and contralateral cognitive areas including the paracingulate and cingulate gyri. This was particularly evident in early windows compared to the sham group (see Figure 2A; Table 2). In later windows both the anodal and cathodal groups had generally reduced variability in connectivity from Lobules I-V to bilateral motor regions, such as the precentral gyrus and the supplemental motor area compared to the sham group. However, during late windows cathodal stimulation increased variability in connectivity between lobules I-IV and cortical regions associated with inhibition and emotion regulation such as the cingulate and paracingulate gyri compared to the sham group (Figure 2B; Table 2). This may suggest that cerebellar stimulation to motor regions differentially impacted variability in connectivity over time post stimulation, which might result in differing behavioral impacts over the course of task completion, when tDCS is used in task paradigms.

**Figure 2.**
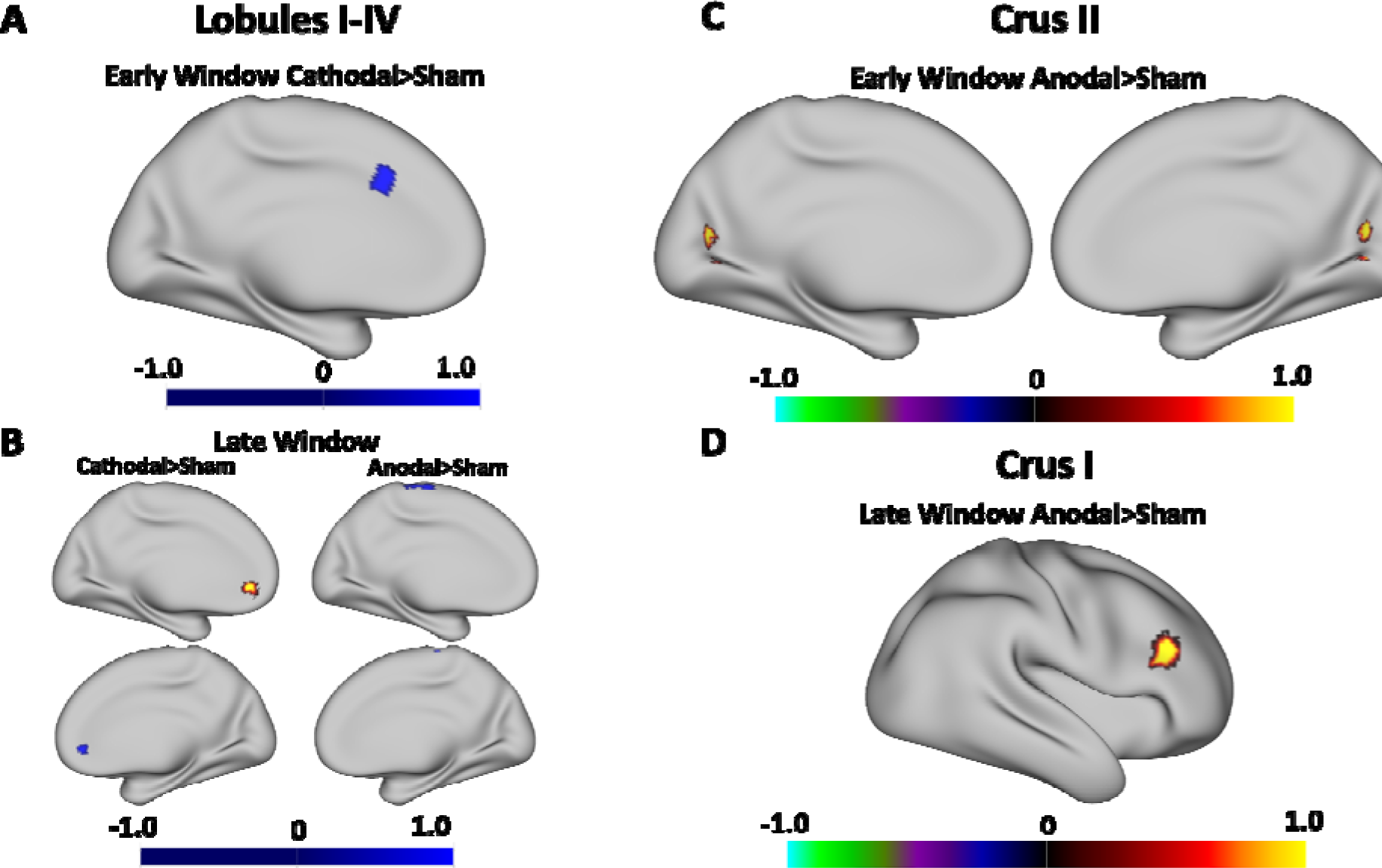
**Left.** The temporal dynamics of connectivity of Lobules I-IV was differentially impacted in the cathodal and anodal stimulation groups relative to the sham group. **A)** During the early window, variability in connectivity was lower in the cathodal group relative to sham. **B)** In the late window both higher and lower variability were observed. Anodal stimulation reduced variability in the late window. **Right.** Connectivity dynamics of both Crus I and II was impacted by anodal stimulation (relative to sham). Variability in both **C)** early and **D)** late windows was higher in regions of the lateral prefrontal cortex and medial visual regions. The color bars denote the percentile range of z-scores. The maps are thresholded such that only significant (p-FDR_J<_J0.05) results are presented.

**Table 2.**
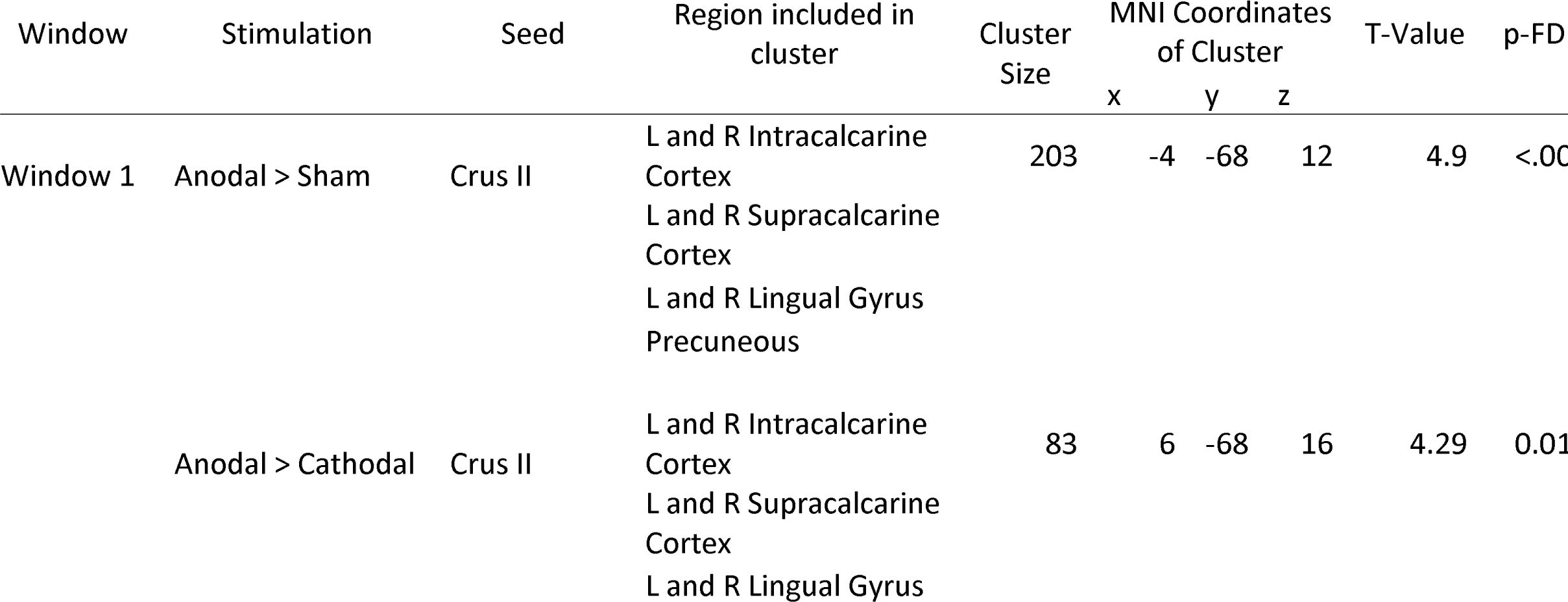
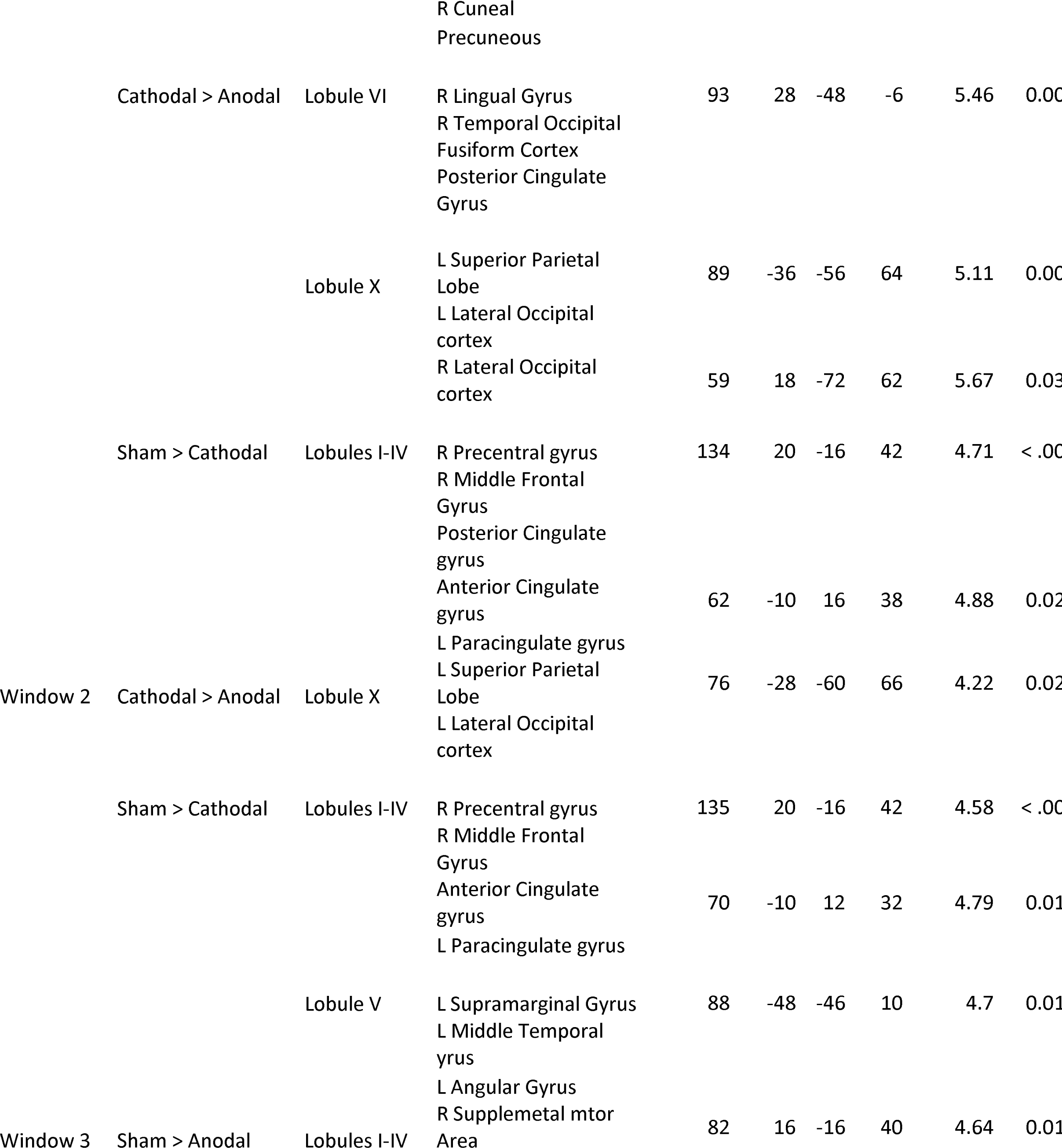
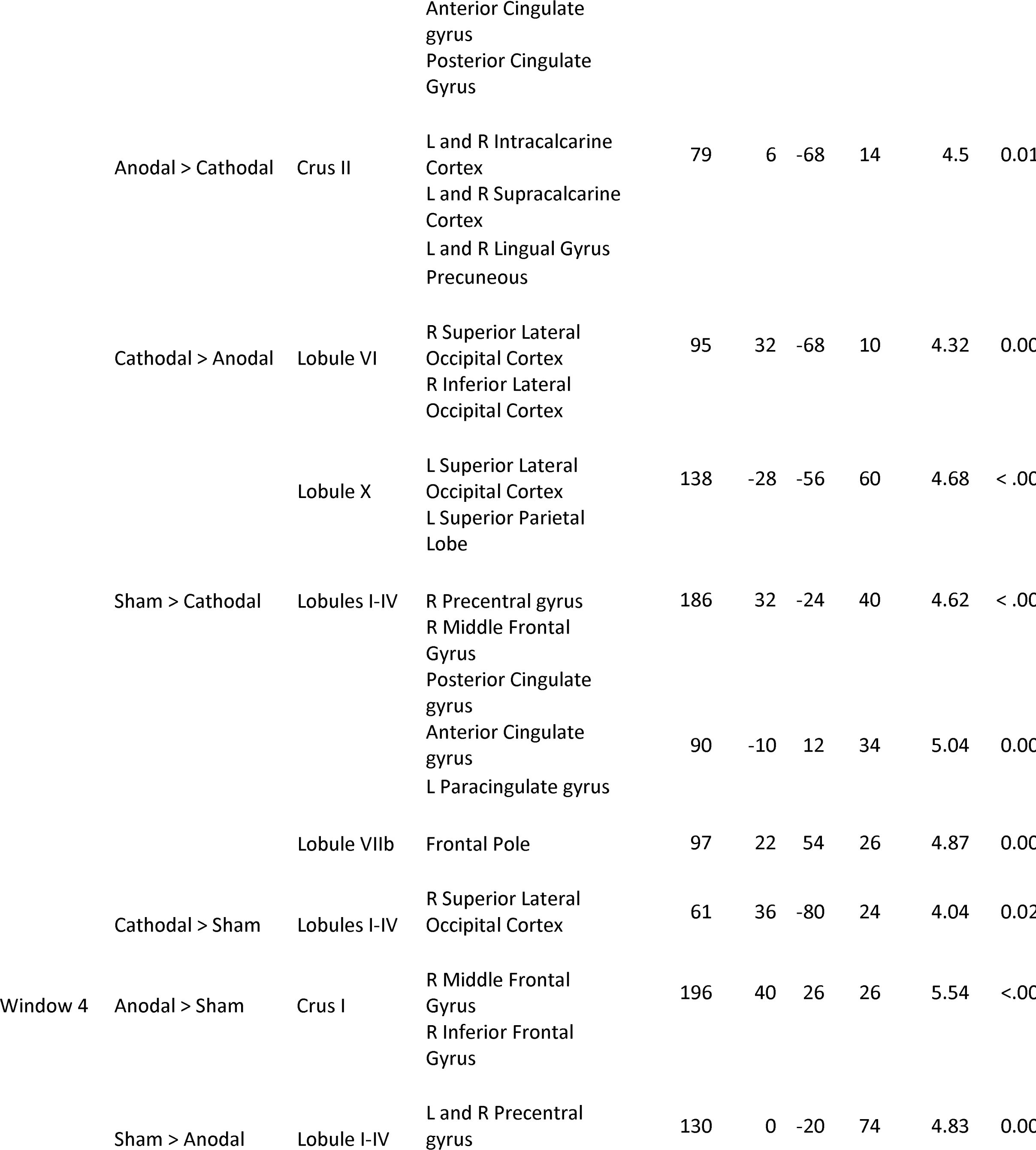
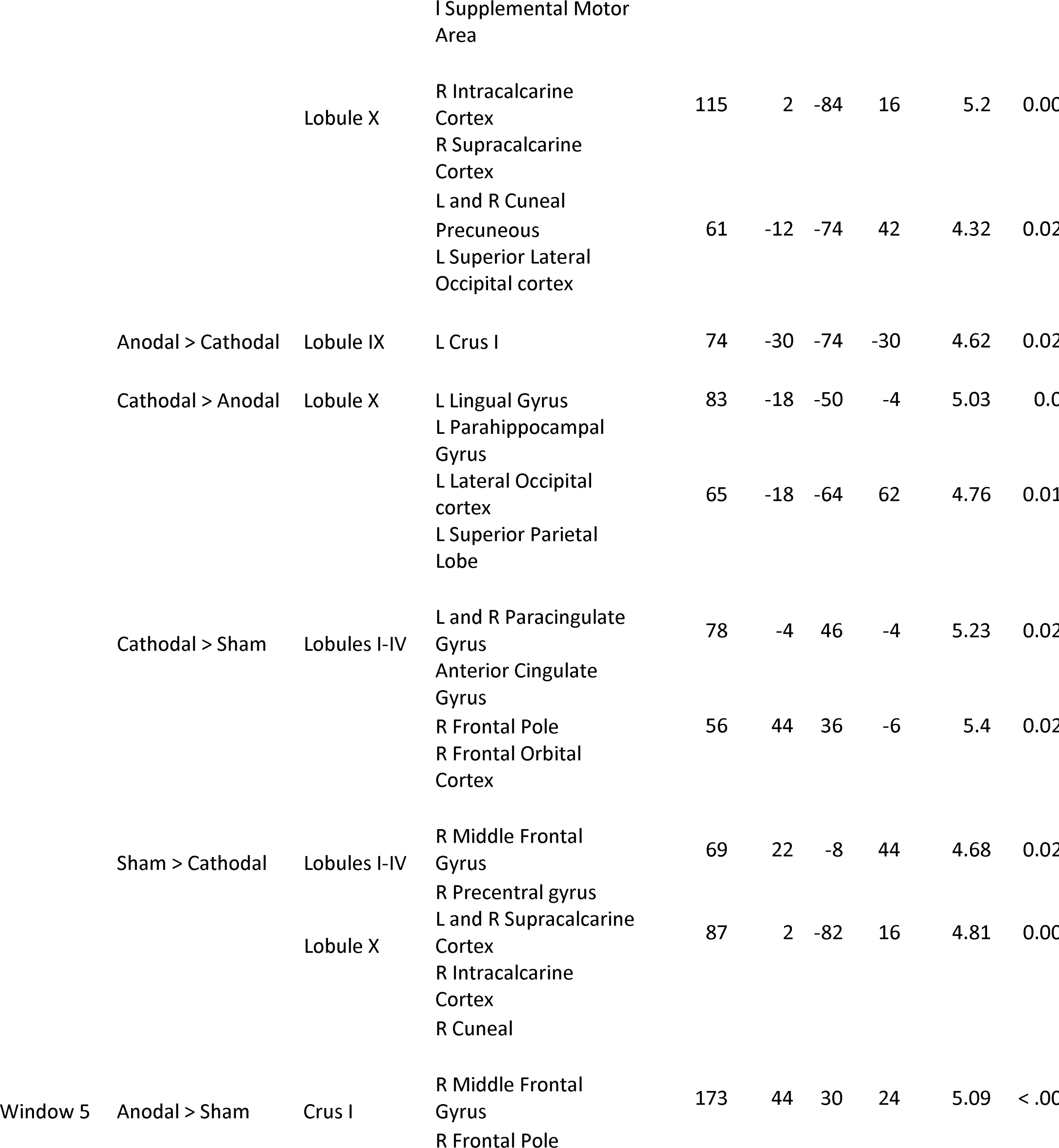
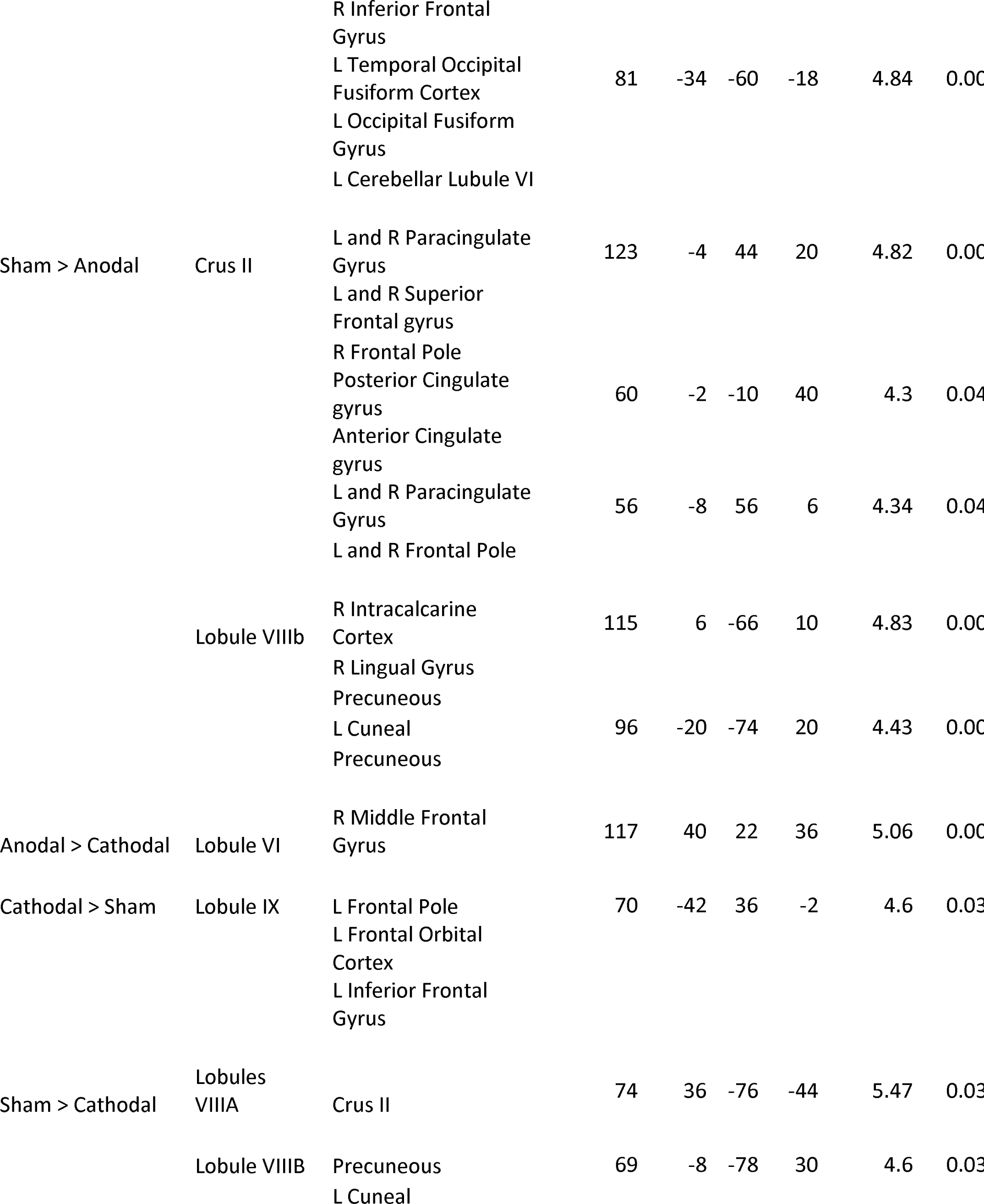
MNI coordinates of the local maxima of brain regions showing functional connectivity with the lobules of the right cerebellar hemisphere by stimulation contrast within each sliding window.

When examining cognitive regions of the right cerebellum, there was a general trend in which anodal stimulation over the right cerebellum resulted in greater variability in connectivity across windows, particularly when analyzing the right Crus I and II seed regions. Much of the variability in connectivity during early windows were found between Crus I and II in the cerebellum and visual regions of the occipital lobe such as the bilateral intracalcarine cortex, supracalcarine cortex, and the precuneus in the anodal stimulation group when compared to the sham and cathodal stimulation group (Figure 2C; Table 2). During later windows, however, variability in connectivity from right Crus I in the anodal stimulation group was greater in language processing centers in the ipsilateral middle and inferior frontal gyrus compared to the sham group (Figure 2D; Table 2). Interestingly, there was less variability in connectivity in the anodal stimulation group between Crus II and cognitive centers in the frontal lobe, compared to sham (Figure 2D; Table 2). This continues to suggest a time dependent nature to cerebellar tDCS.

Additionally, in more posterior cerebellar lobules, such as right lobules VIII-X, the anodal stimulation group had lower variability in connectivity to visual attention centers in the contralateral superior parietal lobule and the lateral occipital cortex compared to the cathodal stimulation group (Table 2). This was particularly evident in early windows in lobule X in the anodal stimulation group. In later windows, both the anodal and cathodal stimulation group saw lower variability in connectivity to visual regions of the occipital lobe such as the bilateral intracalcarine cortex, supracalcarine cortex compared to sham. Interestingly, in late windows, variability in connectivity between lobule IX and X and frontal regions was larger in the cathodal stimulation group to compared to sham (Table 2). This continues to suggest a time dependent nature to the cortical impacts of cerebellar tDCS, particularly when influencing cognitive networks between the cerebellum and cortex.

## Discussion

Cerebellar tDCS can be used to understand how cerebellar networks with the cortex differ after perturbation, which can in turn provide insights into aging as well as neurological and psychiatric illnesses where the cerebellum and cerebello-cortical networks are impacted (Ferrucci et al., 2016; van Dun & Manto, 2018). To this point, however, most work has focused on language networks, with little work looking at other nonmotor processing domains using a whole brain approach. The current work used tDCS over the right cerebellum to better understand cerebello-cortical connectivity. We found stimulation primarily affected cerebellar connectivity to motor and vision processing centers, with some modulation of cortical regions primarily implicated in cognition. Generally anodal stimulation resulted in increased connectivity in both ipsilateral and contralateral regions of the cortex, perhaps indicative of a compensatory response to degraded cerebellar output. A sliding window analysis also demonstrated a time dependent nature to the impacts of cerebellar tDCS on connectivity. Thi was particularly notable for connectivity with cognitive and associative areas of the cerebral cortex. Together, the data suggest the effect of stimulation might be time dependent for both motor and cognitive processing, such that behavioral change because of stimulation might differ depending on when behavior is observed compared to stimulation offset. A better understanding of how temporal dynamics of connectivity changes across time might help interpret behavioral effects of cerebellar stimulation. Our findings are discussed in more detail below.

### Connectivity Patterns and Stimulation

*Motor Regions.* Broadly, when examining connectivity in the sham stimulation group we found the cerebello-cortical connectivity patterns one might to expect between the cerebellum and motor regions in the cortex (Buckner et al., 2011; King et al., 2019). When comparing stimulation groups, connectivity between the cerebellum and motor regions (e.g., precentral gyrus and supplemental motor area) was lower in the cathodal group compared to sham. A sliding window analysis demonstrated that the cathodal stimulation group had lower variability in connectivity to motor regions in the cortex compared to the sham group, during early windows. In late windows, the cathodal stimulation group had increased variability in connectivity to inhibitory centers (paracingulate gyrus) and the anodal stimulation group had decreased variability in connectivity to motor regions compared to sham. Critically, both the cathodal and anodal stimulation group generally saw lower connectivity relative to sham. We predicted only the anodal stimulation group would experience lower connectivity (Galea et al., 2009; Grimaldi et al., 2016), though variability in how cerebellar tDCS affects behavior is not uncommon, particularly when examining motor function (Buch et al., 2017). As such, this mixed literature also would support higher connectivity. Continued work is needed to determine why this variability exists, though our work here may provide some initial insights.

First, stimulation parameters, such as electrode placement and size might not be conducive to modifying activations in lobules I-V in the cerebellum (Rampersad et al., 2014). Indeed, investigation of electric field distributions in the cerebellum suggest anterior regions of the cerebellum, such as Lobules I-V, and the Vermis are not well stimulated (Rampersad et al., 2014). This is the result of the curvature of the cerebellum and its relative anatomical location below the brain. Further, recent work suggested that stimulating different cerebellar regions can have different behavioral and neural effects (Rice et al., 2021). Specifically, connectivity between lobule VIIIb and the cortex in the anodal stimulation group increased, but there was no effect when examining connectivity from lobule V, when compared to the sham group. This further provides evidence that motor lobules of the cerebellum are difficult to target and modulate and electrode placement is crucial in inducing neural and behavioral change.

Second, the cerebellum is a complex structure and it is difficult to know how it might be impacted by stimulation, as the effect depends on current flow direction, relative to axonal position (Chan & Nicholson, 1986). This coupled with the curvature of the cerebellum may contribute to the variable effects seen here and elsewhere (Buch et al., 2017). This lends itself to the ongoing discussion about individual differences in the responsiveness of tDCS (Labruna et al., 2019). Third, it is still not clear which cells types are actually influenced by cerebellar tDCS (Grimaldi et al., 2016). Our hypotheses are based on the modulation of Purkinjie fibers, but their activity is not readily captured by fMRI, instead climbing fibers might drive the fMRI signal (Grimaldi et al., 2016). Though optogenetic work suggests Purkinjie cells respond to cerebellar tDCS (Grimaldi et al., 2016), it is unclear whether other cells are also modulated. Though these findings are not particularly consistent with past work, this does provide further data that indicates how cerebellar tDCS might affect cerebello-cortical connectivity, specifically that the impact of stimulation may vary based on region and timing of stimulation relative to stimulation offset.

*Non-motor regions.* Stimulation to nonmotor regions resulted in significant connectivity differences when compared to the sham stimulation group, but this connectivity was primarily to visual processing regions in the occipital lobe and language centers in the frontal lobes. Critically, our sliding window analysis demonstrated that modulation of connectivity may be time dependent, such that variability of connectivity to language centers was not evident until a later time window. These results were partially in line with past work from Galea et al. (2009) that suggest that anodal stimulation would increase and cathodal stimulation would decrease cortical connectivity. This is also consistent with past cerebellar tDCS and imaging work that found that anodal tDCS to the right cerebellum increased Crus I and II activation and connectivity in cortical language centers (D’Mello et al., 2017) and areas involved in the motor control of speech (Turkeltaub et al., 2016). However, this is the first study to demonstrate impacts to frontal, temporal, and occipital regions associated with cognitive processing outside of language regions as a result of cerebellar stimulation.

The general pattern of results seems to suggest that anodal stimulation gives rise to increased connectivity with ipsilateral regions in the cortex, possibly suggesting a greater need for cortical scaffolding. That is, down regulation in the cerebellum following anodal stimulation might result in the need for greater connectivity to the ipsilateral hemisphere to compensate for the degraded communication between the cerebellum and contralateral cortical regions. As this did not consistently occur in the cathodal stimulation group, we suggest that the degraded output of the cerebellum is driving what we argue may be compensatory connectivity. Interestingly, optogenetic work has suggested that anodal stimulation would negatively impact activation to the cortex (Galea et al., 2009; Grimaldi et al., 2016). Perhaps this impact is not widespread, but hemisphere dependent. That is, decreased contralateral cortical connectivity resulted in the ipsilateral cortex upregulating cortical processing, which might have implications during functional task performance (Maldonado, et al., 2022).

Another notable effect was the time dependent nature of stimulation over the right cerebellum. Previous work has strongly relied on behavioral data (Boehringer et al., 2013; Ferrucci et al., 2008; Majidi et al., 2017; Maldonado et al., 2019; Pope & Miall, 2012; Spielmann et al., 2017; van Wessel et al., 2016; Verhage et al., 2017) with tDCS, but the results have typically been mixed. The time dependent nature of cerebellar tDCS could explain the mixed nature of the cerebellar tDCS literature. That is, differences in behavioral outcomes might be dependent on when the effect of cerebellar tDCS is realized in the cortex. This time dependent nature might also have specific methodological implications, such that timing between cerebellar tDCS and task completion might be a factor to consider when constructing stimulation procedures. Currently, the behavioral literature is largely mixed (Oldrati & Schutter, 2018), but predicts polarity specific outcomes, regardless of stimulation parameters or task domain (Galea et al., 2009), such as task completion relative to stimulation offset. However, it is possible that modulation of specific outcome variables might be dependent on specific stimulation parameters, as evidenced by differences in connectivity depending on time and domain.

Rather unexpectantly, cathodal stimulation did not seem to meaningfully impact non-motor connectivity to the cortex when compared to the anodal and sham stimulation groups. This is not in line with our hypothesis, though past work focused on language processing has demonstrated increased connectivity and behavioral performance following cathodal stimulation (Pope & Miall, 2012; Turkeltaub et al., 2016). Critically, this work, to our knowledge, is the first attempt to assess the effect of cathodal stimulation over the right cerebellum on cerebello-cortical connectivity broadly.

Here we suggest that in this sample of healthy younger adults, there are polarity dependent group differences in cerebellar frontal interactions with stimulation that support the idea of degraded cerebellar processing and communication with the cortex after stimulation. Research in aging and disease might provide insights into degraded cerebello-cortical interactions. Specifically, disruptions to cerebellar function and network connectivity in aging (Bernard et al., 2013) or disease (Allen et al., 2007; Bai et al., 2009) may negatively impact prefrontal function and processing. Cerebellar resources might be important for supporting cortical processing and provide crucial scaffolding for normative performance and function. However, anodal stimulation might disrupt this scaffolding by perturbing cerebello-cortical connectivity, specifically the temporal dynamics of connectivity. That is, the not only is the degree of connectivity important, but also the time course of the interaction. Existing compensatory scaffolding models broadly suggest increased activation to compensate for lost ability in advanced age (Cabeza, 2002; Cabeza, Anderson, Locantore, & McIntosh, 2002; Park & Reuter-Lorenz, 2009; Reuter-Lorenz & Campbell, 2008; Reuter-Lorenz & Park, 2014), but these models focus primarily on the prefrontal cortex and general degradation of functional ability, though past work suggests that the cerebellum might be involved (Bernard et al., 2013; Maldonado, et al., 2022). The current work might parallel the perturbed cerebello-cortical connectivity found in advanced age or disease, which would suggest some cerebellar contribution to cortical processing, allowing a better understanding of cerebello-cortical interactions and how these interactions support cortical scaffolding. The inclusion of the cerebellum in these compensatory scaffolding models might help explain the mixed effects of cerebellar tDCS on behavior, particularly when accounting for the degree to which connectivity is affected and the time-dependent nature of this effect, in relation to cortical processing broadly.

## Conclusions

The current work aimed to better understand the impact of cerebellar tDCS on cerebello-cortical connectivity. Stimulation group differences in connectivity provided novel and important insights into the time dependent effects of tDCS on cerebellar networks. This work provides new insights into cerebello-cortical connectivity in the face of perturbation which has implications for aging and both neurological and psychiatric disease. Specifically, initial support for the cerebellum as a component for existing scaffolding models might be evident in the current data, further supporting its inclusion in models describing frontal lobe activation in health and disease. However, at this point additional work that further considers tDCS stimulation parameters and functional boundaries is still warranted.

## Acknowledgements

We would like to thank research assistants Sydney Eakin, Ivan Herrejon, and Sydney Cox for their help with data collection. Portions of this research were conducted with the advanced computing resources provided by Texas A&M High Performance Research Computing.

## Declarations

JAB was supported in part by R01 AG064010-01. The authors have no relevant financial or non-financial interests to disclose. The authors do not have any conflicts of interest. The data that support the findings of this study are available from the corresponding author upon reasonable request.

## Notes

### Competing Interest Statement

The authors have declared no competing interest.

